## Supplementary Materials for "KM-GPT: An Automated Pipeline for Reconstructing Individual Patient Data from Kaplan–Meier Plots"

#### A Supplementary Figures and Tables

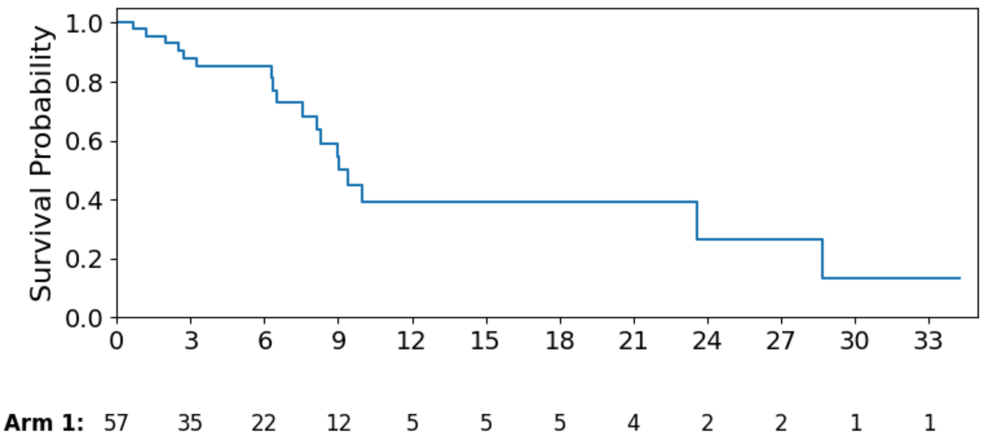


(A) LLH

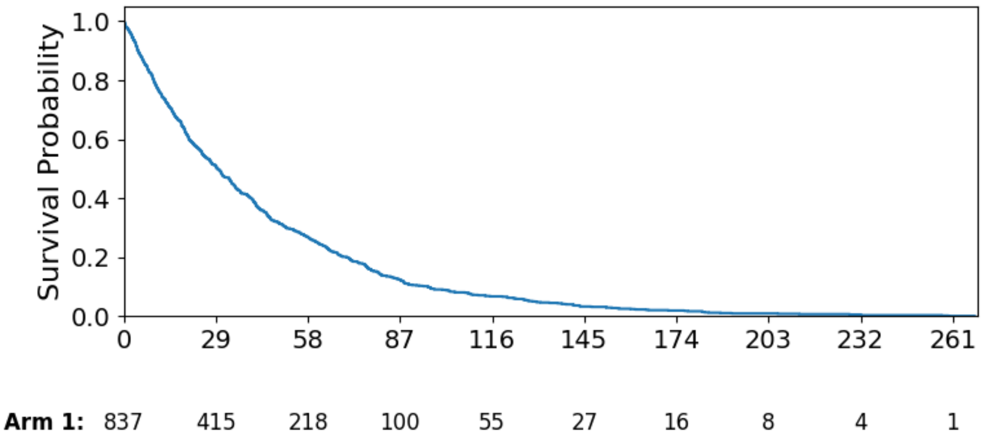


(B) HHL

Figure S1: Examples of Simulated Kaplan–Meier Survival Curves. Simulation groups (e.g., LLH and HHL) denote combinations of study sample size, median survival time, and censoring rate, with each letter representing a low (L), medium (M), or high (H) setting for the corresponding parameter.

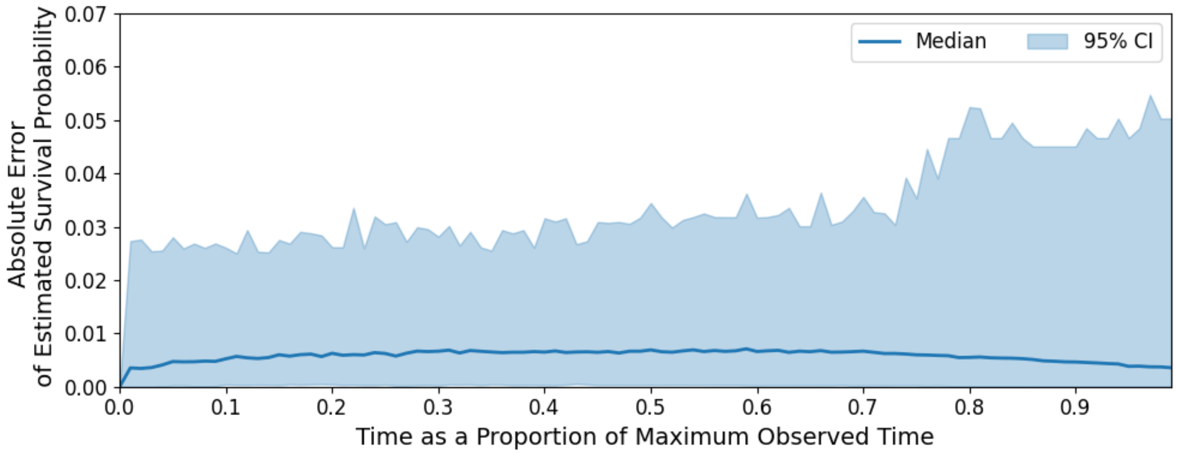


Figure S2: Absolute Error between Reconstructed Survival Curves and Ground Truth across Normalized Time.

Table S1: Comparison of Reported and Reconstructed mOS in PD-L1 CPS Subgroups.

| Trial | Treatment | Reported mOS | Reconstructed mOS |
|---|---|---|---|
| **PD-L1 CPS** $\geq 1$ | | | |
| KEYNOTE-061 | Pembrolizumab | 9.1 (6.2 – 10.7) | 9.06 (6.23 – 10.98) |
| KEYNOTE-062 | Pembrolizumab | 10.6 (7.7 – 13.8) | 10.72 (8.40 – 14.28) |
| JAVELIN Gastric 100 | Avelumab | 14.9 (8.7 – 17.3) | 14.97 (8.32 – 17.26) |
| **PD-L1 CPS** $\geq 10$ | | | |
| KEYNOTE-061 | Pembrolizumab | 10.4 (5.9 – 18.3) | 10.75 (6.11 – 17.77) |
| KEYNOTE-062 | Pembrolizumab | 17.4 (9.1 – 23.1) | 17.52 (9.08 – 22.12) |
| JAVELIN Gastric 100 | Avelumab | 8.2 (3.9 – NR) | 6.31 (3.94 – 18.00) |

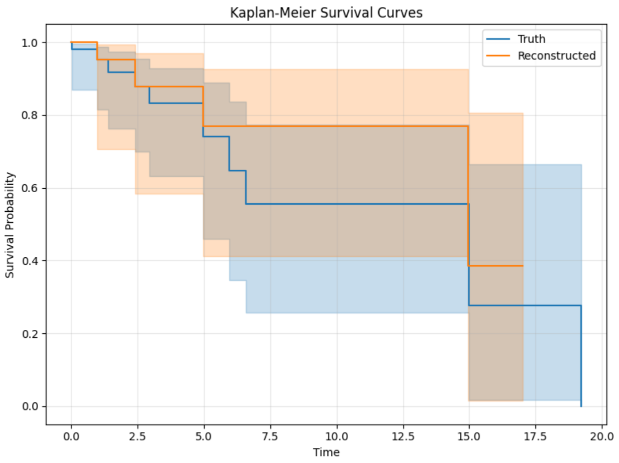


(A) LLH

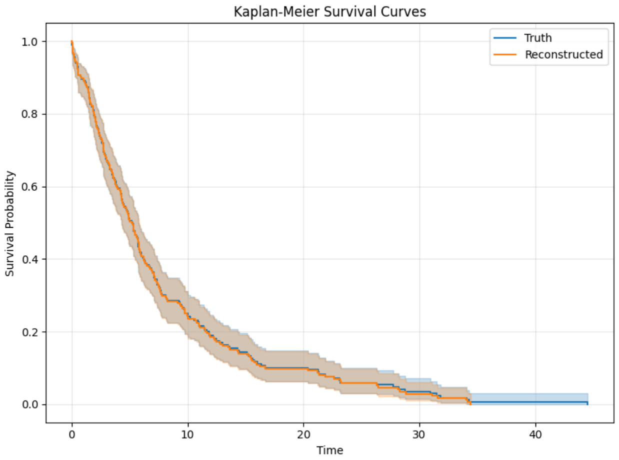


(B) MHL

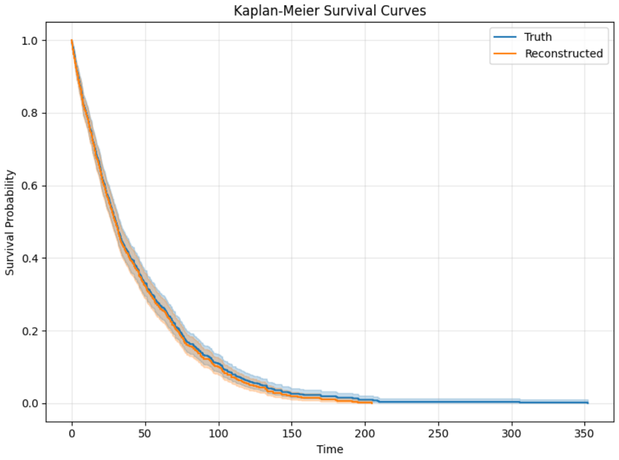


(C) HHL

Figure S3: Representative Examples of Poor Reconstructions from Low-Performance Groups. Panel A (LLH) shows a small-sample, high-censoring scenario where steep drops and sparse events in the later follow-up period result in unstable curve estimation. Panels B (MHL) and C (HHL) illustrate a large study with a low censoring rate, where the final portion of the curve is truncated.

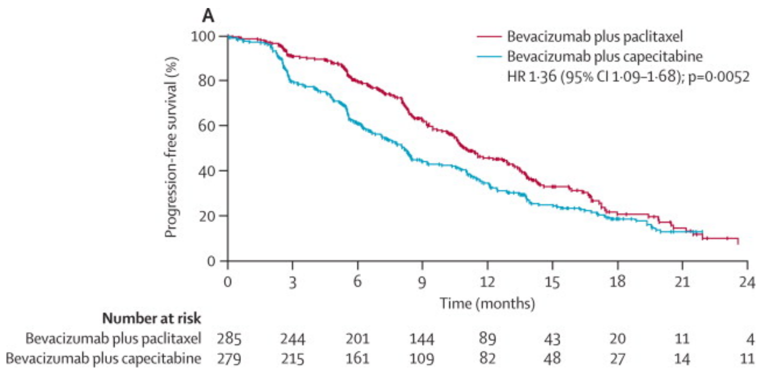


(A) PMID 23312888

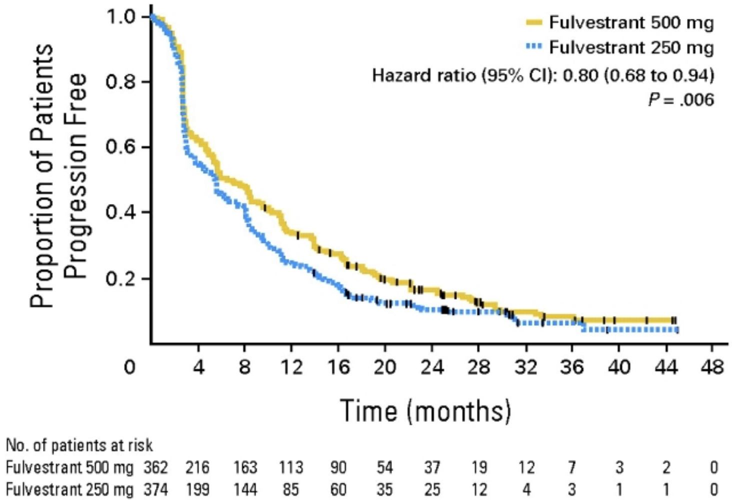


(B) PMID 20855825

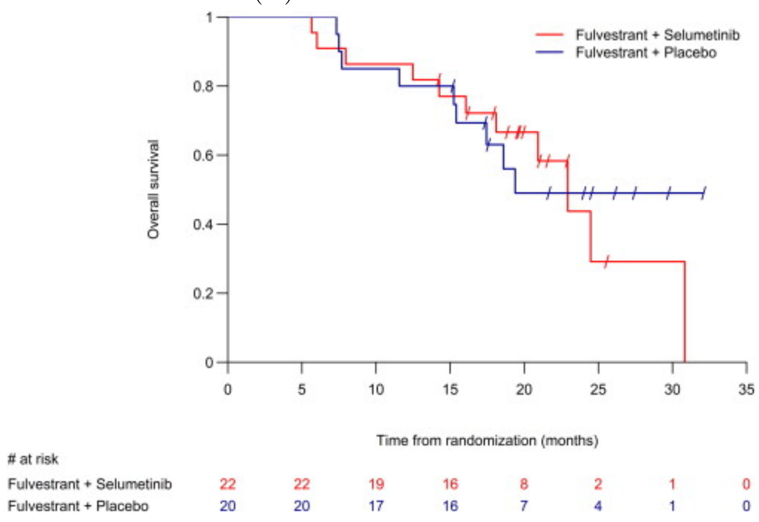


(C) PMID 25892646

Figure S4: Original KM Plots from Three Metastatic Breast Cancer Studies.

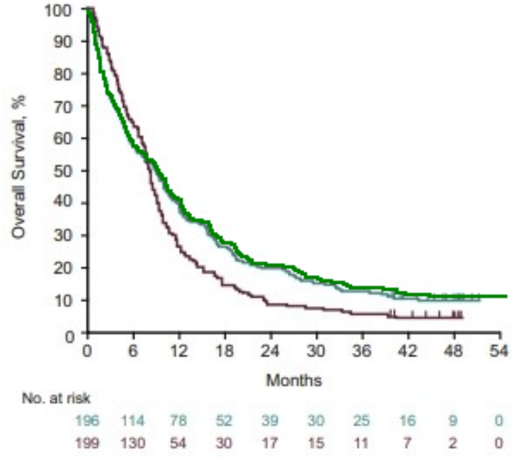


(A) KEYNOTE-061 (CPS $\geq$ 1)

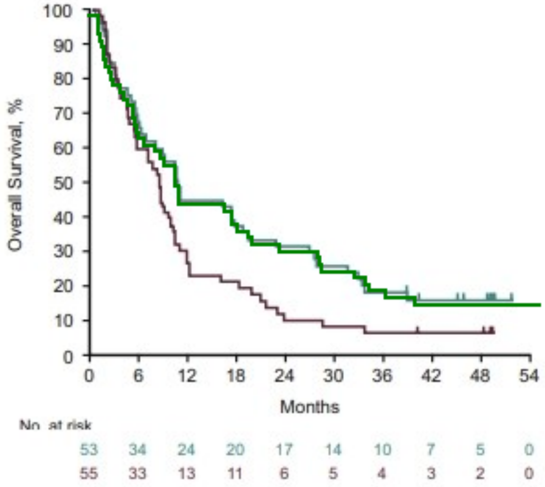


(B) KEYNOTE-061 (CPS $\geq$ 10)

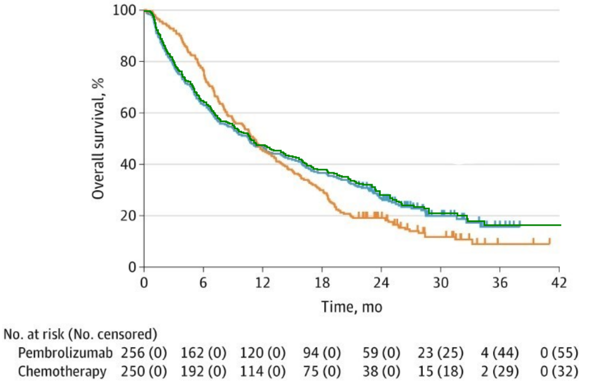


(C) KEYNOTE-062 (CPS $\geq$ 1)

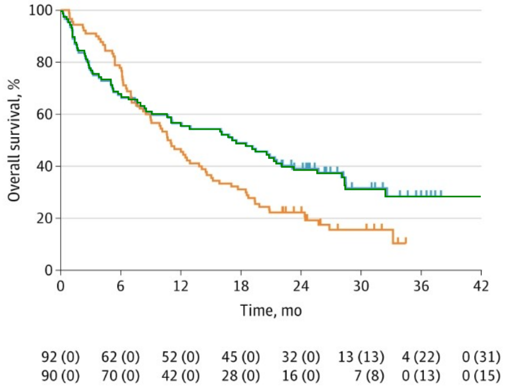


(D) KEYNOTE-062 (CPS $\geq$ 10)

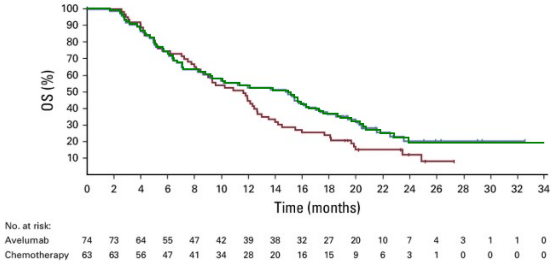


(E) JAVELIN Gastric 100 (CPS $\geq$ 1)

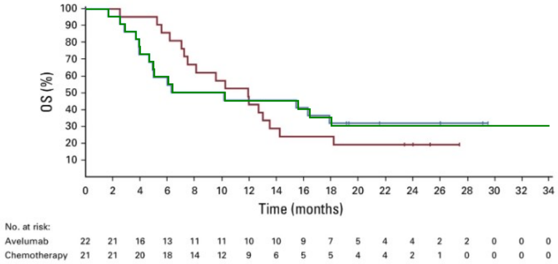


(F) JAVELIN Gastric 100 (CPS $\geq$ 10)

Figure S5: Original KM Plots from Trials with Reconstructed KM Overlaid. Green lines are the reconstructed KM curves from KM-GPT overlaid on ICI treatment arms from original figures.

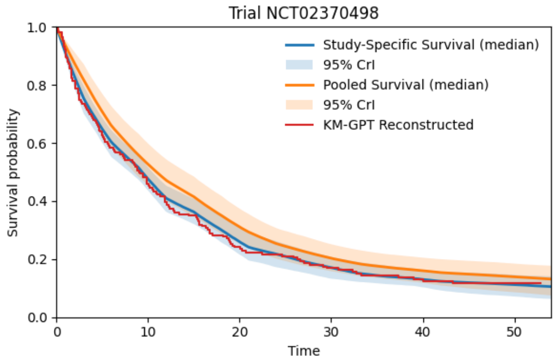


(a)

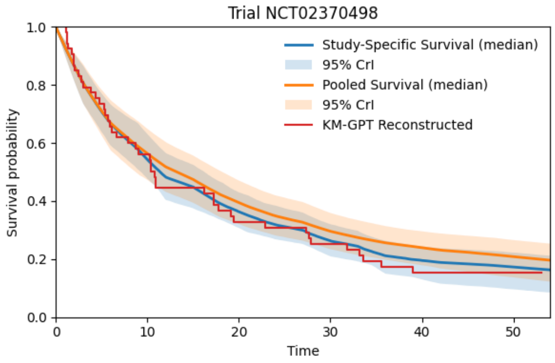


(b)

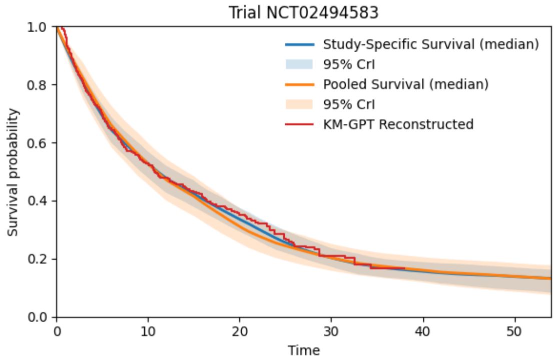


(c)

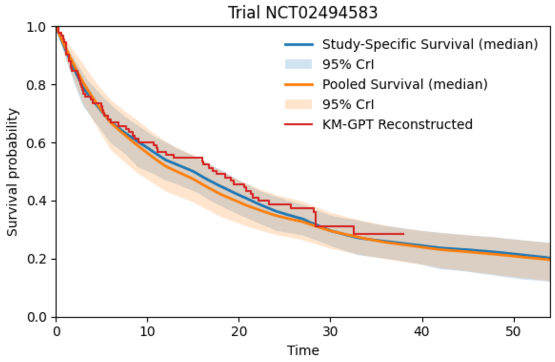


(d)

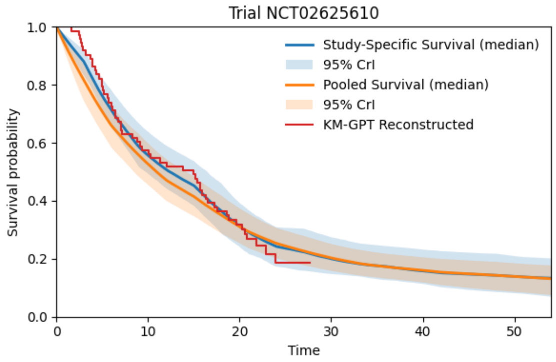


(e)

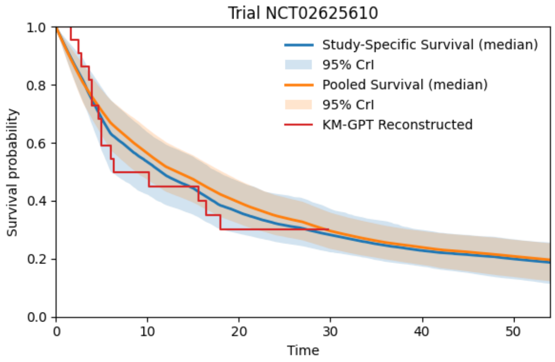


(f)

Figure S6: Posterior Inference of Study-Specific and Pooled Survival Curves.

### B OCR Engine Settings

To optimize text extraction from Kaplan–Meier plots, we configured the OCR engine with the following parameters:

- **OCR Engine Mode:** KM-GPT utilizes the **oem 3** mode of the OCR engine, which leverages an advanced LSTM-based model for text recognition. This mode is highly accurate across a variety of fonts and image qualities, ensuring reliable text extraction.
- **Risk Table Loading Mode**: For loading risk tables, we set the engine to **psm 6**, a mode specifically designed for extracting structured content, such as rows and columns, from number-at-risk tables.

Combining **oem 3** for high character recognition accuracy and **psm 6** for structured data extraction ensures the robust and consistent parsing of layout-dependent textual information in Kaplan–Meier plots. These preprocessing steps form the foundation for the Multi-Modality Processing Unit (MMPU), which further refines and interprets OCR outputs using GPT-5 to achieve high-fidelity table reconstruction.

### C KM-GPT Technical Details

#### C.1 Axis Calibration

Following preprocessing, axis calibration is performed to map image pixels into real-valued time and survival probability domains. Tick labels extracted via OCR are parsed by the *range detection* routine, which infers four key parameters: $t_{\min}, t_{\max}, s_{\min}, s_{\max}$. Unique numeric labels are then sorted, and pairwise gaps between labels are computed. The most common increment is identified from a trimmed histogram of these differences, forming the time increment $\Delta_t$ and survival probability increment $\Delta_s$. For irregular or non-monotonic sequences, the median of the increments is used as the $\Delta$ value.

Axis endpoints and orientations are determined by the axis identification procedure, which projects ink densities along columns and rows of the lightness channel to locate vertical and horizontal axis strokes. From these projections, the pixel coordinates of the axis baselines $(u_{x_0}, u_{x_1})$ and $(v_{y_0}, v_{y_1})$ are established, and interior margins are automatically adjusted based on surrounding whitespace gradients.

Finally, an affine transformation is applied to convert pixel coordinates $(u, v)$ into real-world values using the equations:

$$t(u) = t_{\min} + \frac{u - u_{x_0}}{u_{x_1} - u_{x_0}} (t_{\max} - t_{\min}), \qquad s(v) = s_{\max} - \frac{v - v_{y_0}}{v_{y_1} - v_{y_0}} (s_{\max} - s_{\min}),$$

where the inversion of $s(v)$ accounts for the image origin being in the top-left corner. This calibration ensures sub-pixel accuracy in quantifying digitized survival curves. The same transformations are also used to convert curve pixels into their corresponding time and survival probabilities.

#### C.2 Curves Differentiation

To partition foreground pixels into curve-specific groups, we use a color-space partitioning approach. For each pixel, features $(h, s, l)$ in HSL color space are extracted and stored in a DataFrame. Near-background pixels are optionally removed by retaining only those with lightness $l \geq 0.2$, effectively suppressing pale grid-lines and page artifacts. The features are then standardized for clustering: if image enhancement is applied, the $s$ (saturation) and $l$ (lightness) channels are up-weighted by a factor of 100 to amplify chroma and luminance differences relative to the $h$ channel. We fit a $K$-medoids model [29] with $K =$ Num of Curves using Euclidean distance in $(h, s, l)$. The best model, determined by minimizing inertia, is selected, and pixels are assigned to clusters based on the nearest medoids, producing distinct curve labels. The result is a grouped pixel set, where each color represents one unique curve.

#### C.3 Overlapping Curve Interpolating

To address overlapping segments on curves, we implement a local $k$-NN consensus scoring method followed by a grid-constrained path tracing technique. Pixels are first sorted and embedded in a Euclidean $k$-NN graph [30], where the labels of neighboring pixels are analyzed. For each pair of neighbors $i$ and $j$, we construct a same-group indicator matrix $\mathbb{I}_{ij} \in \{-1, +1\}$, where $+1$ indicates the neighbor $j$ shares $i$'s cluster label and $-1$ otherwise. Neighbor contributions are weighted using inverse-squared distance, $w_{ij} = 1/(d_{ij}^2 + \varepsilon)$, where $\varepsilon = 10^{-10}$ ensures numerical stability for very small separations. The per-pixel consensus score, estimating local label confidence in overlapping or parallel regions, is computed as the normalized weighted sum: $\text{score}_i = \frac{1}{k} \sum_j \mathbb{I}_{ij}\, w_{ij}$. Using these consensus scores, we trace the curves and interpolate points in the overlapping regions, ensuring accurate segment reconstruction where curves overlap or run closely together.

### D Hierarchical Piecewise-Exponential Model for Meta Analysis

#### D.1 Model Formulation

In this section, we present the Bayesian hierarchical model for meta-analysis. The time axis is partitioned into $J$ disjoint intervals, $I_j = (t_{j-1}, t_j]$ for $j = 1, \ldots, J$, within which the hazard function for each study is assumed to be constant.

The hazard rate for study $s$ in interval $j$ is $\lambda_{sj} = \exp(\alpha_{sj})$, where $\alpha_{sj}$ is the study- and interval-specific log-hazard. To pool information across studies, each $\alpha_{sj}$ is modeled as a Gaussian deviation from a pooled, interval-specific log-hazard $a_j$:

$$\alpha_{sj} \mid a_j, \sigma_j^2 \sim \mathcal{N}(a_j, \sigma_j^2). \tag{D.1}$$

Here, $a_j$ represents the overall meta-analytic log-hazard in interval $j$, and $\sigma_j^2$ captures the between-study heterogeneity for that interval.

To share strength across adjacent time intervals and ensure the pooled hazard evolves smoothly, we place a hierarchical prior on the sequence of pooled parameters $\mathbf{a} = (a_1, \ldots, a_J)$. Specifically, we model $a_j$ as varying around a latent mean $\mu_j$:

$$a_j \mid \mu_j, \sigma_a^2 \sim \mathcal{N}(\mu_j, \sigma_a^2), \tag{D.2}$$

where the latent process $\boldsymbol{\mu} = (\mu_1, \ldots, \mu_J)$ follows a stationary autoregressive process of order one (AR(1)) to enforce smoothness:

$$\begin{aligned} \mu_1 \mid \phi, \tau^2 &\sim \mathcal{N}\left(0, \frac{\tau^2}{1-\phi^2}\right), \\ \mu_j \mid \mu_{j-1}, \phi, \tau^2 &\sim \mathcal{N}\left(\phi\mu_{j-1}, \tau^2\right), \quad \text{for } j = 2, \ldots, J, \end{aligned}$$

with $|\phi| < 1$ ensuring stationarity and $\tau > 0$.

We complete the model specification with the following prior distributions. The standard deviation parameters $\sigma_j$ (characterizing between-study heterogeneity), $\sigma_a$ (governing variation of the pooled effects around the latent mean), and $\tau$ (the innovation standard deviation of the latent process), are assigned weakly informative Half-Normal priors: $\sigma_j \sim \mathcal{N}^+(0, 0.2^2)$, $\sigma_a \sim \mathcal{N}^+(0, 0.2^2)$, and $\tau \sim \mathcal{N}^+(0, 1^2)$. The autoregressive parameter $\phi$ is modeled via a transformed parameter to maintain the stationarity constraint $|\phi| < 1$; specifically, we define $\phi = \tanh(\psi)$ and assign the prior $\psi \sim \mathcal{N}(0, 0.75^2)$.

The pooled hazard in interval $j$ is $\lambda_j^{\text{pool}} = \exp(a_j)$, and for study $s$ the study-specific hazard is $\lambda_{sj} = \exp(\alpha_{sj})$. The corresponding survival functions at time $t$ are derived from the cumulative hazard. Let $\Delta_k = t_k - t_{k-1}$ be the length of the $k$-th interval and let $j(t)$ be the index of the interval such that $t \in I_{j(t)}$. The survival probability for the pooled population is given by:

$$S_{\text{pool}}(t) = \exp\left(-\sum_{k=1}^{j(t)} \lambda_k^{\text{pool}} \Delta_k\right),$$

and for study $s$, it is given by:

$$S_s(t) = \exp\left(-\sum_{k=1}^{j(t)} \lambda_{sk} \Delta_k\right).$$

Figure S6 displays the posterior survival trajectories from the hierarchical piecewise-exponential model. Each panel illustrates the reconstructed survival curve, the study-specific curves, and the pooled survival function. Shaded bands represent 95% credible intervals, derived from 5000 posterior draws after 2000 warm-up steps of Markov Chain Monte Carlo (MCMC) sampling. The study-specific curves capture heterogeneity across trials, reflecting variability in population size, follow-up duration, and censoring patterns. In contrast, the pooled curve provides a stable summary that smooths out trial-level fluctuations. These results demonstrate the model's ability to recover individual trial survival patterns while also producing a coherent pooled estimate that balances study-specific evidence with cross-study information.
